# Flagella are required to coordinately activate competition and host colonization factors in response to a mechanical signal

**DOI:** 10.1101/2023.12.31.573711

**Authors:** Lauren Speare, Liang Zhao, Morgan N. Pavelsky, Aundre Jackson, Stephanie Smith, Bhavyaa Tyagi, Garrett C. Sharpe, Madison Woo, Lizzie Satkowiak, Trinity Bolton, Scott M. Gifford, Alecia N. Septer

**Affiliations:** Department of Earth, Marine & Environmental Sciences, University of North Carolina, Chapel Hill, NC; Department of Microbiology, Oregon State University, Corvallis, OR; Department of Chemistry, Morgan State University, Baltimore, MD

**Keywords:** *Aliivibrio fischeri*, viscosity, transcriptomics, type VI secretion, aggregation

## Abstract

Bacteria employ antagonistic strategies to eliminate competitors of an ecological niche. Contact-dependent mechanisms, such as the type VI secretion system (T6SS), are prevalent in host-associated bacteria, yet we know relatively little about how T6SS+ strains make contact with competitors in highly viscous environments, such as host mucus. To better understand how cells respond to and contact one another in such environments, we performed a genome-wide transposon mutant screen of the T6SS-wielding beneficial bacterial symbiont, *Vibrio fischeri*, and identified two sets of genes that are conditionally required for killing. LPS/capsule and flagellar-associated genes do not affect T6SS directly and are therefore not required for interbacterial killing when cell contact is forced yet are necessary for killing in high-viscosity liquid (hydrogel) where cell-cell contact must be biologically mediated. Quantitative transcriptomics revealed that *V. fischeri* significantly increases expression of both T6SS genes and cell surface modification factors upon transition from low-to high-viscosity media. Consistent with coincubation and fluorescence microscopy data, flagella are not required for T6SS expression in hydrogel. However, flagella play a key role in responding to the physical environment by promoting expression of the surface modification genes identified in our screen, as well as additional functional pathways important for host colonization including uptake of host-relevant iron and carbon sources, and nitric oxide detoxification enzymes. Our findings suggest that flagella may act as a mechanosensor for *V. fischeri* to coordinately activate competitive strategies and host colonization factors, underscoring the significance of the physical environment in directing complex bacterial behaviors.

**Significance:** The physical environment can have dramatic effects on bacterial behavior, but little is known about how mechanical signals impact antagonistic interactions. Symbiotic bacteria use molecular weapons to eliminate competitors for limited space within highly viscous host tissue and mucus.

To better understand how the physical environment affects competition and adhesion within eukaryotic hosts, we used quantitative transcriptomics to reveal the flagella-dependent transcriptional response to bacterial transition from lower to a higher viscosity environment. This work revealed the T6SS interbacterial weapon is coordinately activated with host colonization factors, emphasizing the importance of integrating activation of interbacterial weapons into host colonization pathways to enhance a symbiont’s ability to successfully colonize the host while efficiently eliminating potential competitors from the host niche.

## Introduction

Bacterial competition is prevalent in free-living and host-associated communities and a myriad of competitive strategies have been identified (1). Many of these mechanisms require direct cell-cell contact for inhibitor cells to deliver effector proteins directly into target cells (2). On surfaces, bacteria often grow in dense communities that facilitate this necessary contact. However, little is known about how bacteria make contact in liquid environments. In such environments contact-dependent mechanisms may be particularly useful as diffusible antimicrobial molecules would become quickly diluted. The type VI secretion system (T6SS) is a contact-dependent mechanism that allows inhibitor cells to deliver effector proteins directly into target cells through a syringe-like apparatus. T6SSs are prevalent in free-living, symbiotic, and pathogenic species (3, 4) and are active in animal microbiomes (5–10).

The bioluminescent squid and fish symbiont *Vibrio fischeri* encodes a strain-specific T6SS (T6SS2) that allows cells to eliminate competitors during symbiosis establishment (5, 11). *V. fischeri* activates T6SS2 expression and killing upon transition from a low-to high-viscosity liquid environment, which mimics the increase in viscosity cells experience during host colonization (12). In high-viscosity liquid (hydrogel) *V. fischeri* make large, mixed-strain aggregates that are biologically mediated and pH-dependent; these aggregates facilitate the contact necessary for T6SS-mediated killing in a liquid environment (12, 13). We recently identified a putative cell-cell adhesion gene associated with T6SS2, *tasL,* that is required for target specificity and inhibitor-target contact in hydrogel via a heterotypic interaction with an unknown ligand (11). In addition to viscosity, the presence of calcium is also sufficient to activate T6SS2 expression and aggregation in low-viscosity liquid (14). However, the way in which *V. fischeri* cells sense and respond to changes in environmental viscosity to activate T6SS2 and promote cell-cell contact requires further investigation.

Previous work in other model organisms has revealed several ways in which a bacterial cell can sense changes in mechanical signals within its physical environment (15), including both flagella-dependent and independent mechanisms. Flagella-dependent mechanisms rely on sensing a change in torque in the motor when flagella experience an increase in resistance from encountering a surface or more viscous fluid. Flagella-independent mechanisms include external sensory proteins such as pilins, or integral membrane proteins that sense pressure changes. Sometimes these mechanisms are used in combination, where flagella propel the cell through a liquid environment and an external sensor detects the change in the environment.

*V. fischeri* genomes encode multiple proteins with possible connections to mechanosensory abilities, including flagella, pilins, and membrane-associated proteins with predicted mechanosensitive domains. The lophotrichous polar flagella are a collection of multiple flagella that are sheathed by the outer membrane and anchored at one point, acting in concert to drive motility in a single direction (16, 17). In the context of symbiosis, flagella provide propulsion to migrate along the colonization route to the light organ crypts and provide a key mechanism for signaling to the host via outer membrane vesicle (OMV) and flagellin release (18–21). Although previous data suggests *V. fischeri* is capable of mechanosensing (12, 13), which we define here as a biological response to a mechanical cue (i.e. viscosity) (22), the genetic determinants of *V. fischeri* mechanosensory mechanisms remain unknown.

Here, we sought to identify such factors by performing an unbiased screen to identify genes required for killing in high-viscosity liquid conditions. We screened transposon mutants of the T6SS2-encoding *V. fischeri* strain MJ11 and identified mutants that cannot kill a target in hydrogel but are still lethal on surfaces, suggesting they are unable to kill under conditions that require biologically mediated cell-cell contact. We characterized the impact of these “conditional” mutations, including flagellar mutants, on T6SS sheath assembly, aggregation, and motility. We then used quantitative transcriptomics to explore the role of flagella in regulating gene expression during habitat transition. The quantitative transcriptomic data generated from this analysis reveals the significant transcriptional shift *V. fischeri* cells undergo during habitat transition, the contribution of flagella in responding to changes in environmental viscosity, and insight into how a model beneficial bacterial symbiont coordinates activation of key competitive and colonization pathways in response to its physical environment.

## Results

### Transposon mutagenesis of MJ11 identifies genes required for killing in hydrogel

To begin identifying genetic factors required for target cell elimination in hydrogel, we generated transposon mutants of the lethal *Vibrio fischeri* strain MJ11 and then screened these mutants for the ability to prevent the growth of an unarmed target strain, ES114, in hydrogel and on surfaces. MJ11 encodes the strain-specific T6SS2 genomic island (5) and kills ES114 in a T6SS2-dependent manner (13). MJ11 has a fully sequenced genome, allowing for identification of transposon insertions throughout the genome (23), and importantly is brightly luminescent in culture, compared to ES114 (24), which allowed us to discriminate between MJ11 and GFP-tagged ES114 in coculture. We incubated GFP-tagged ES114 with wild-type MJ11 or a T6SS structural mutant (*tssF_2/vasA_2*) as controls for target killing and target survival, respectively (Fig 1A). Briefly, each MJ11-derived strain was mixed with GFP-tagged ES114 at a 9:1 (MJ11:ES114) ratio by volume and incubated in hydrogel (liquid LBS + 5% polyvinylpyrrolidone [PVP]) without shaking for 24 hours. To screen many mutants simultaneously, coincubation assays were optimized for 96-well plates, as described previously (11), and competitive outcomes were determined by the relative fluorescence units (RFU) of the GFP-tagged target strain. Any mutant that was unable to reduce the target RFUs was then confirmed through additional coincubation assays in hydrogel where target strain colony forming units (CFUs) were quantified at the beginning and end of each experiment (Fig 1B) (12). We screened 7,300 mutants and identified 20 mutants that could no longer kill ES114 in hydrogel (Fig 1B and Table S1) but could still prevent the growth of GFP-tagged ES114 when cell-cell contact was forced on agar surfaces (Fig 1C).

**Figure 1.**
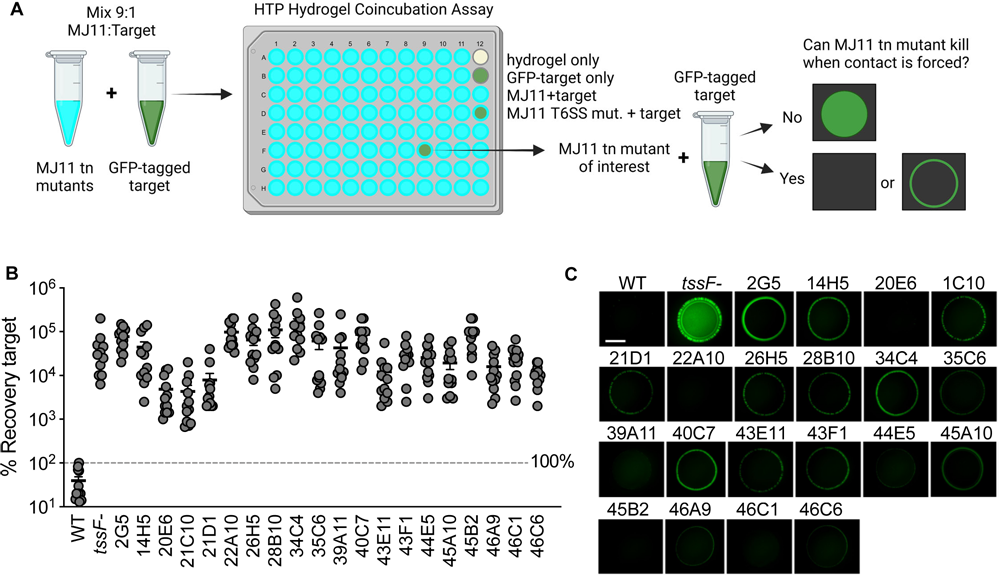
Transposon screen identified 21 conditional mutants. (A) Approach for initial screen to identify nonlethal MJ11 Tn mutants in hydrogel (made with Biorender.com). Tn mutants were arrayed into a 96-well plate containing LBS hydrogel medium and a culture of GFP-tagged ES114 and cultures were mixed in a 9:1 MJ11:ES114 ratio. After 24-hours, each well was assessed for luminescence and GFP to determine the presence of MJ11 and ES114, respectively. Wells with luminescence and fluorescence values similar to the target + MJ11 T6SS-control well were selected for future experiments. (B) Results of coincubation assays in hydrogel between MJ11 wild-type (WT), T6SS2 disruption mutant (*tssF-*) or transposon mutant and ES114 (target) displayed as percent recovery of the target strain. MJ11 strain genotype is listed on the x-axis; transposon mutant names are abbreviated as follows: LAS2G5 shown as 2G5. Dashed line at 100% recovery indicates no net change in target growth over the 12-hour coincubation period. (C) Results of coincubation assays on agar surfaces between MJ11 strains and ES114 shown as fluorescence microscopy images of ES114 taken at 24 hours; scale bar = 2 mm. MJ11 strain name is shown above each coincubation spot. Each experiment was performed three times and either combined data (B, n=12) or a representative replicate are shown (C, n=1).

We used a combination of cloning and inverse PCR to determine the transposon insertion site for each conditional MJ11 mutant. The transposons disrupted 12 unique genes found only on chromosome I (Table S1). The majority of disrupted genes were in a predicted capsule/LPS biosynthesis gene cluster (9 genes). Transposons also disrupted flagellar motility genes (*flgK* and *flgB*) and a putative deaminase/amidohydrolase. Given that conditional genes are not required for killing on surfaces, we hypothesized that the conditional genes identified here are involved in mediating the necessary cell-cell contact for T6SS killing in hydrogel.

### Conditional genes are not required for T6SS2 sheath assembly

To begin characterizing the role of conditional genes in T6SS-mediated competition, we tested whether any conditional genes were necessary for constructing a T6SS sheath. VipA/TssB is a subunit of the T6SS sheath and by tagging it with GFP, we can visualize sheath assembly in live cells (5). We moved a VipA_2-GFP expression vector into a representative transposon mutant for each disrupted conditional gene and used fluorescence microscopy to visualize sheath assembly. Wild-type MJ11 cells contained fully extended GFP-tagged sheaths, while the MJ11 *tssF_2* mutant control strain, which lacks the essential baseplate component, contained no sheaths and only diffuse GFP was observed in these cells (Fig 2A). All conditional mutants contained VipA_2-GFP sheaths (Fig S1). This observation suggests that conditional genes are not necessary for T6SS2 sheath assembly and is consistent with our hypothesis that conditional genes are involved in mediating the cell-cell contact required for killing in hydrogel.

**Figure 2.**
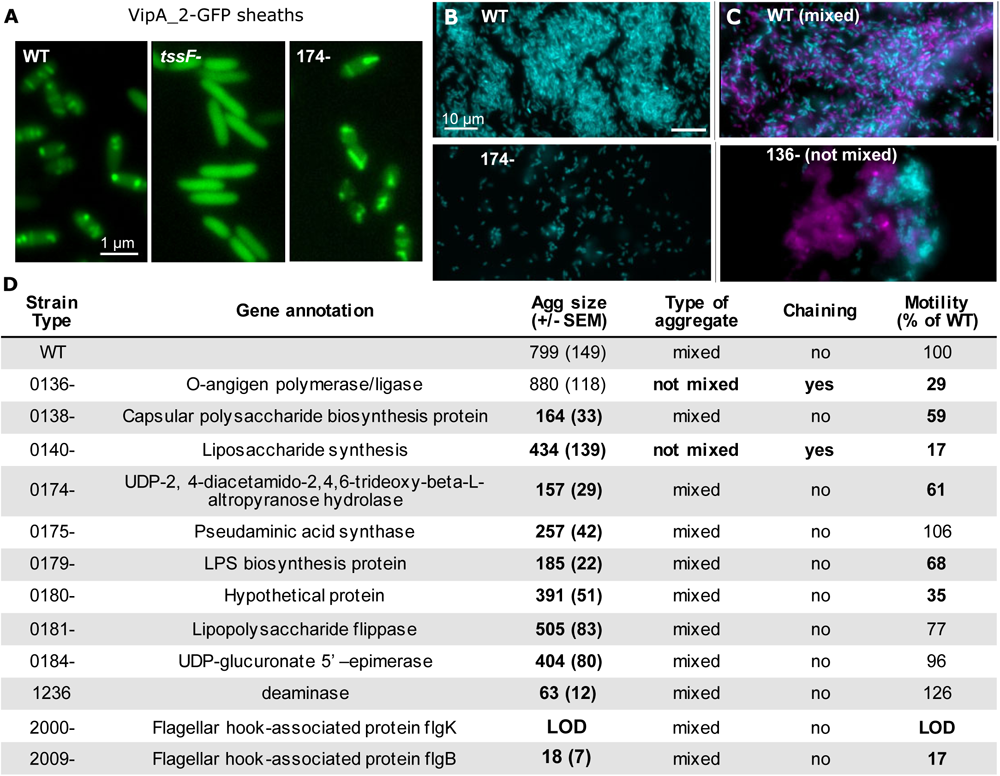
Motility and LPS / capsule genes are necessary for mixed-strain aggregates in hydrogel. (A) Representative fluorescence microscopy images of MJ11 WT, *tssF-,* or 174-harboring an IPTG-inducible VipA_2-GFP expression vector. Cultures were incubated in hydrogel supplemented with 1.0 mM IPTG; scale bar = 1 um. Sheaths were observed in all transposon mutants. Each experiment was performed twice with 2 biological replicates and 2-3 fields of view (n=10). (B & C) Representative single-cell fluorescent microscopy images of (B) MJ11 monocultures or (C) differentially-tagged but otherwise isogenic MJ11 co-cultures incubated in hydrogel for 12 hours; scale bar = 10 um. (D) Table displays results from experiments with representative transposon mutants where column 1 indicates the disrupted gene and column 2 shows the annotation for each gene. Columns 3-6 display results from the following experiments: 3) monoculture assays quantifying aggregation (Agg) size after 12 hours in hydrogel; 4) co-culture assays with differentially-tagged but otherwise isogenic MJ11 strains to determine whether aggregates were evenly mixed (C, top) or not mixed (C, bottom); 5) visual observation of whether cells displayed chaining in hydrogel; and 6) motility assays in LBS motility agar reported as percent of the wild-type. Each experiment was performed at least twice with two biological replicates and five fields of view per replicate (Aggregation assays, Chaining assays; n=20) or three times with four biological replicates (Motility assay; n=12). SEM indicates standard error of the mean; LOD indicates limit of detection.

### Surface modification and motility mutants are impaired in aggregation in hydrogel

The majority of disrupted conditional genes fall into two predicted functional categories: surface modification (putative capsule/LPS biosynthesis genes) and flagellar motility (*flgK* and *flgB)* (Table S1). To test the hypothesis that conditional genes are required for the necessary cell-cell contact for T6SS2 killing in hydrogel, we assessed the ability of representative conditional mutants (one mutant per disrupted gene) to form wild-type sized mixed-strain aggregates, because we previously showed that deficiencies in either of these factors would reduce cell-cell contact between inhibitor and target and result in impaired T6SS-mediated killing in hydrogel (11, 13). The majority of transposon mutants (11 out of 12) had a significantly smaller average estimated aggregate size compared to the wild type (Fig 2B and 2D, Fig S2). 10 of these 11 aggregation-deficient mutants formed mixed-strain aggregates with differentially-tagged yet otherwise isogenic strains, with both strain types distributed throughout the aggregates (Fig 2C, 2D, S3). Two mutants (genes VFMJ11_0136 and VFMJ11_0140), however, formed aggregates that were not well mixed and were composed of disparate clusters of each strain type (Fig 2C and S3). In total, we observed deficiencies in either the ability to form wild-type sized and/or mixed strain aggregates in all 12 of the transposon mutants we tested. Together, these observations are consistent with our hypothesis that the genes identified in our transposon screen are required for wild-type like aggregation, and therefore necessary for T6SS2 function in hydrogel.

### Motility is necessary but not sufficient for aggregate formation

Given that the two flagellar motility mutants were unable to form substantial aggregates (Fig 2D), we predicted that motility was required for aggregate formation in hydrogel. To test this prediction, we performed motility assays in hydrogel media supplemented with 0.3% agar to quantify motility of each strain at the population level. As expected, the two flagellar mutants had significantly reduced motility relative to the wild type and the *flgK* mutant did not display any motility during the experiment (Fig 2D). The majority of conditional mutant strains tested also had significantly reduced motility compared to the wild type (Fig 2D). However, three surface modification mutants had wildtype levels of motility yet made significantly smaller aggregates, suggesting that motility is required, yet not sufficient to facilitate wildtype aggregation in hydrogel.

### *flgK* is required for transcriptional up-shift during habitat transition

Given that flagella play a key role in environmental sensing in other bacteria (15, 25) and are required in *V. fischeri* for aggregation in hydrogel (Fig 2), we sought to investigate the regulatory role of flagella during habitat transition. Scanning electron microscopy (SEM) confirmed the *flgK* mutant does not produce polar flagella (Fig 3A) and genome resequencing determined no secondary mutations in genes related to aggregation factors (Fig S4). Moreover, when the *flgK* transposon mutation was moved into a fresh MJ11 background using natural transformation (26), the remade *flgK* mutant (MP101) showed similar aggregation deficiencies, compared to the original *flgK* mutant (LAS22A10) (Fig S5), indicating that *flgK* is required for proper aggregation.

**Figure 3.**
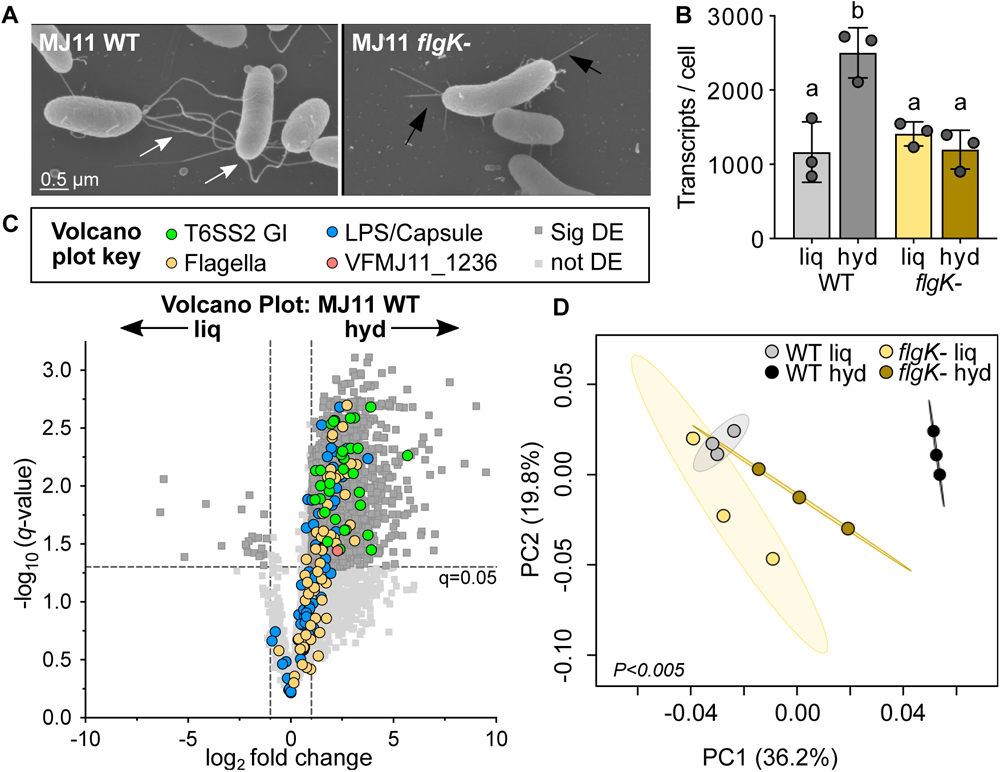
MJ11 gene expression increases dramatically during transition from liquid to hydrogel in a flagella-dependent manner. (A) Scanning electron microscopy (SEM) images of MJ11 wild-type (WT) and *flgK* mutant (*flgK-*) in liquid LBS for 12 hours. Flagella are observed for the wild-type (white arrows) yet not the *flgK* mutant, where pili (black arrows) are still observed. (B) Number of transcripts per cell for MJ11 transcriptomes for the wild-type (WT) or *flgK* mutant (*flgK-*) incubated in liquid LBS (liq) or hydrogel (hyd). Each circle represents an individual sample (transcriptome); letters indicate significantly different values of transcripts per cell between sample groups (two-way analysis of variance [ANOVA] with Šídák’s multiple comparison post-test, *p<0.02*). (C) Volcano plot showing log_2_ fold difference in transcript abundance between liquid and hydrogel for MJ11 wild-type transcriptomes. Each point represents expression of a single gene. Genes with a negative log_2_ fold change have higher expression in liquid (left), while genes with a positive log_2_ fold change have higher expression in hydrogel (right). Colors indicate the functional assignment of genes of interest: green, T6SS2 genomic island (GI); yellow, flagella; blue, LPS/capsule; red, VFMJ11_1236. Data points above the dashed horizontal line had significant q-values between conditions (Student’s t-test with correction for false discovery rate, q<0.05), and those outside the vertical dashed lines had a magnitude fold change of >|1| log_2_ between conditions (dark gray squares). (D) Principal coordinates analysis (PCoA) plot of transcriptome data. The percentages on each axis indicate the amount of variation explained by the axis and ellipses indicate 95% confidence intervals. Adonis P values for comparison between transcriptomes is shown in the lower left-hand corner.

We employed quantitative transcriptomics to achieve two specific aims: 1) determine how habitat transition impacts *V. fischeri* transcript abundance at the per cell level, and 2) describe the role of flagella in mediating the observed changes in gene expression. We modified methods described from Gifford *et al.* (27, 28) to produce quantitative transcriptomes for *V. fischeri* MJ11 wild-type and a flagellar mutant (*flgK-*) identified in our screen. Briefly, RNA was collected from monocultures of each strain grown in liquid or hydrogel for 12 hours, when cells are still in exponential growth (12). Quantitative transcriptomics was achieved by addition of internal RNA standards just prior to RNA extraction (see methods and Text S1 for an in-depth explanation). Internal standard transcript sequences were recovered and used to convert the number of reads per library to number of transcripts per cell, based on CFU values. Moreover, comparing these CFU counts for each treatment revealed no significant difference, suggesting comparable growth and number of cells were used to harvest RNA (Fig S6A).

To obtain a high-level understanding of how cells transcriptionally respond to the physical environment, we first asked whether strain genotype and/or media viscosity affected the amount of mRNA per cell. We evaluated the number of transcripts per cell for each strain in each condition, and observed that on average, wild-type liquid samples had 1,165 transcripts per cell (Fig 3B, S6B), which is comparable to other bacterial species during exponential growth in a lab setting (*E. coli:* 1,380-1800 mRNAs / cell) (29, 30). Unexpectedly, wild-type hydrogel samples had 2,501 transcripts per cell, more than twice as many as in liquid (Fig 3B, S6B). The *flgK* mutant had 1,410 and 1,196 transcripts per cell in liquid and hydrogel, respectively (Fig 3B). The *flgK* mutant mRNA values were not significantly different from the wild type in liquid, but were significantly lower than the wild type in hydrogel (Fig 3B). Thus, the *flgK* mutation prevents an increase in transcription in hydrogel grown cells. We next performed a principal coordinates analysis (PCoA) on the MJ11 transcriptomes and found that *flgK* mutant samples clustered with wild-type liquid samples and separately from wild-type hydrogel samples (Fig 3D ANOVA: *P<0.005*). Taken together, these data suggest that flagella contribute significantly to MJ11’s transcriptional response to increased environmental viscosity.

To identify which transcripts were driving this change in gene expression for wild-type cells, we generated a volcano plot comparing transcript abundance for each gene between liquid and hydrogel (Fig 3C). Approximately 59% of the genome (2,451 genes) was significantly differentially expressed between culture conditions, and most of those genes (2,429 or 99% of differentially expressed genes), had higher expression in hydrogel while only 22 genes (1% of differentially expressed genes) had higher expression in liquid (Fig 3C). This observation suggests that MJ11 upregulates over half of its genome based on a mechanical signal (viscosity) that changes as cells transition from free-living to a host-associated lifestyle. Notable gene transcripts that were significantly more abundant in hydrogel included: i) T6SS2 genes, which we expected based on previous work (12), ii) surface modification and flagellar synthesis gene clusters, which we obtained transposon insertions in for this screen (Fig 3D), iii) 94 predicted transcriptional regulators, and iv) host-relevant metabolic and detoxification pathways. Therefore, we next used hierarchical clustering analysis to determine whether differences in gene expression are driven by condition (liquid vs hydrogel), genotype (WT vs *flgK*-), or a combination of both.

### Flagella promote expression of surface modification genes in hydrogel

Our transcriptomes revealed genes in the surface modification cluster identified in our transposon screen are also significantly upregulated in the wild type in hydrogel. This genomic region, for which we isolated transposon mutants with impaired aggregation and killing, encodes gene products predicted to be involved in LPS modification, synthesis of predicted surface-exposed glycans such as pseudanimic acid, and homologs of components of the Wxz/Wxy capsule biosynthesis and export system (Fig 4A) (31). Hierarchical cluster analysis of transcript abundance for genes in this genomic region revealed that wild-type hydrogel samples clustered separately from all other samples, while wild-type liquid, *flgK* mutant liquid, and *flgK* mutant hydrogel samples clustered together (Fig 4B). On average, wild-type hydrogel samples had higher gene expression compared to the other samples, although this was only statistically significant for 15 of 60 genes (Fig 4B, asterisks). Taken together, these findings support three main conclusions: 1) surface modification genes are generally upregulated in wild-type upon transition from low to high viscosity liquid, 2) this increase in gene expression is dependent on *flgK*, and 3) these surface modification genes are necessary for aggregation required for T6SS-dependent killing in hydrogel.

**Figure 4.**
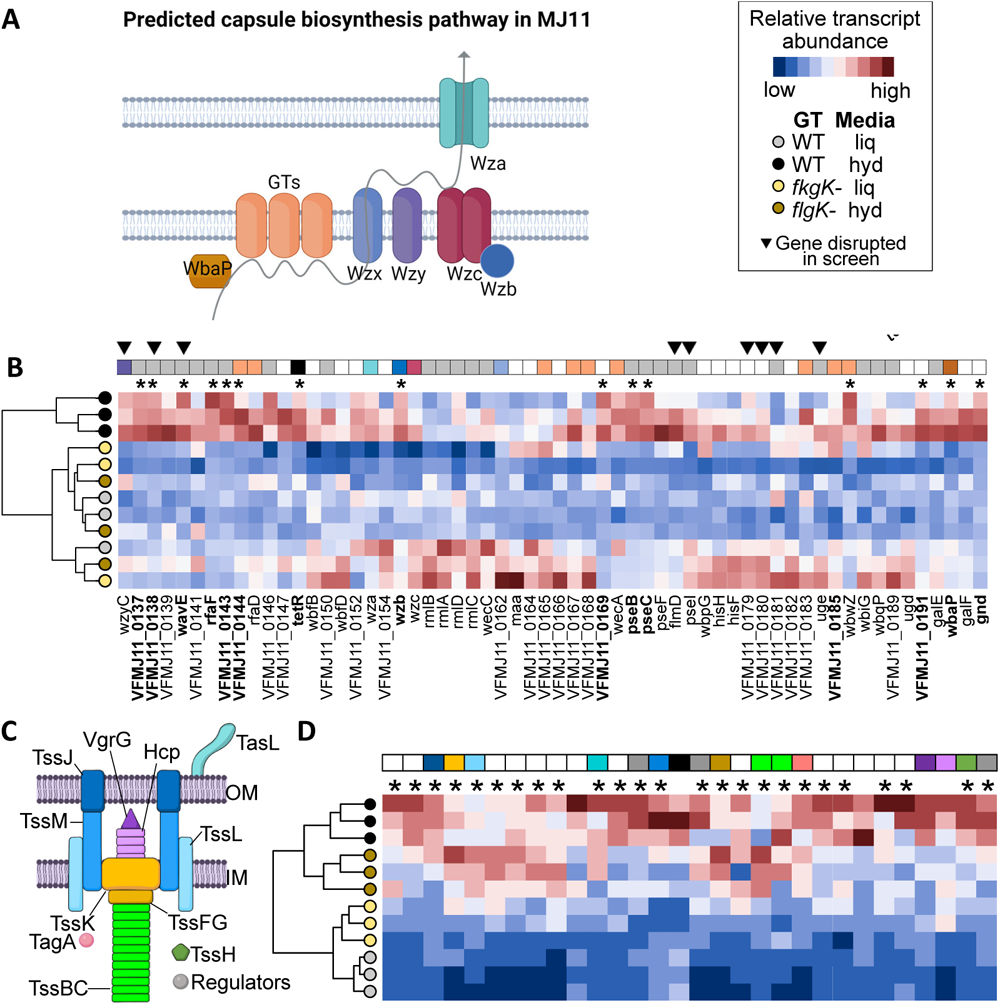
Flagella-dependent gene expression for competition factors in liquid or hydrogel. Hierarchical clustering analysis and heatmap displaying transcript abundance for each gene for the surface modification (A-B) and T6SS2 (C-D) gene clusters. Circle color indicates the strain genotype and media condition: light gray (wild-type [WT] in liquid [liq]), dark gray (wild-type in hydrogel [hyd]), light yellow (*flgK* mutant [*flgK-*] in liquid), and dark yellow (*flgK* mutant in hydrogel). Transcript abundance is shown as relative transcript abundance where values are scaled across each gene (column); blue indicates relatively low transcript abundance while red indicates relatively high abundance. For panels B and D, squares above each column (gene) indicate its known or predicted function based on associated schematic (A and C); OM, outer membrane; IM, inner membrane. Gray boxes indicate predicted polysaccharide synthesis genes. Black boxes indicate predicted transcriptional regulators. For panel B, asterisks indicate expression of a gene was significantly higher for wild-type hydrogel samples compared to all other samples (one-way ANOVA, *p<0.03*). For panel D, asterisks indicate expression of a gene was significantly higher for hydrogel samples compared to liquid samples (Student’s t-test, *p<0.01*). Gene ID (VFMJ11_#) for each heatmap are shown left to right as follows: A0804 – A0833. Figure made with Biorender.com.

Finally, a hierarchical cluster analysis of the T6SS2 gene cluster grouped samples by media condition rather than strain genotype (Fig 4DE). This observation indicates that strain genotype did not influence sample clustering, which is consistent with our above findings that the flagellar components *flgK* and *flgB* are not required for T6SS2 sheath assembly or function on agar surfaces. Together, these data indicate that flagella are not required for T6SS2 activation in hydrogel or on surfaces, suggesting the T6SS weapon is activated by a different sensory mechanism that is coordinated with flagella-dependent expression of predicted surface modification genes. Indeed, pilins were still observed in the *flgK* SEM images (Fig 3A), suggesting that additional mechanosensory mechanisms require exploration. Because *V. fischeri* genomes encode multiple pilin structures, it is possible that our screen would not have identified these, or other mechanical sensors that may have functional redundancy.

### Coculture of *flgK* and *tasL* mutants enhances aggregation and killing in hydrogel

Given the importance of surface modifications described above, and the previous work showing the putative lipoprotein TasL is required for a heterotypic interaction for aggregation in hydrogel, we wondered whether one or more of these surface modifications may interact with TasL. We hypothesized that if TasL and a *flgK*-dependent surface modification interact, then mixing a *tasL* mutant with a *flgK* mutant would enhance aggregation and killing ability in hydrogel, compared to the single mutations, because aggregation and killing ability might be restored if each genotype provides one part of the heterotypic interaction.

To determine whether mixing *flgK* and *tasL* mutants enhances aggregation ability, we coincubated differentially-tagged strains in hydrogel and imaged aggregates to determine the relative size of aggregates and whether mixed-strain aggregates would form. Control treatments with differentially-tagged MJ11 parent, the *tasL* mutant, or the *flgK* mutants yielded the expected results. MJ11 formed large, mixed strain aggregates while the *tasL* mutant formed small, mixed strain aggregates, and the *flgK* mutant had no aggregation (Fig 5A). However, when we mixed the *flgK* and *tasL* mutants, we saw both strain types present in larger aggregates. (Fig 5A).

**Figure 5.**
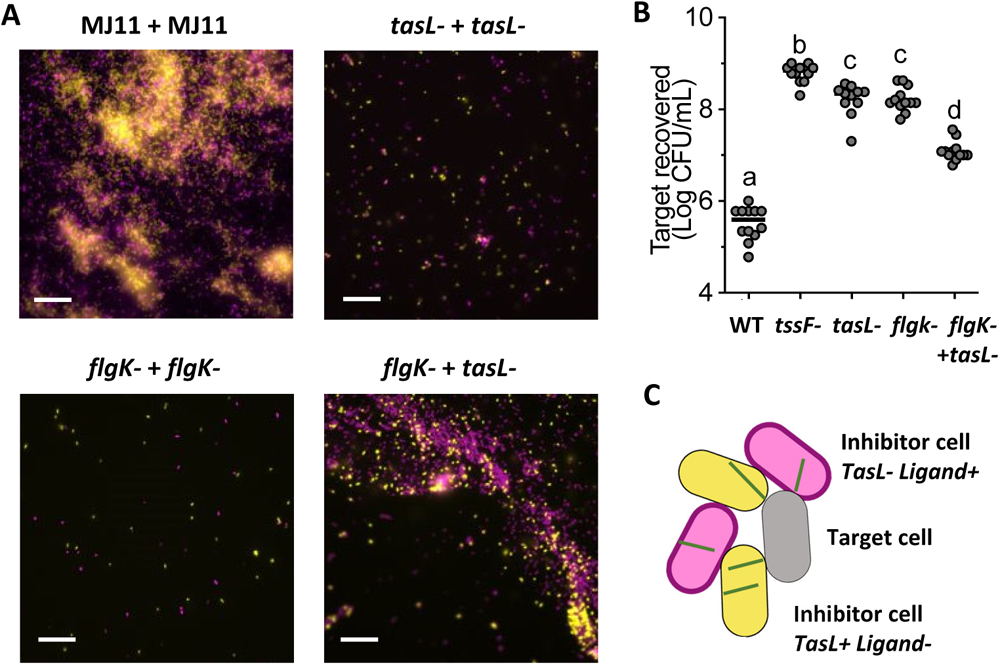
*flgK* and *tasL* mutant coculture restores aggregation and killing in hydrogel. (A) Representative images of coincubation of differentially-tagged strains in hydrogel. Scale bar is 20 µm. (B) Results from coincubation assays between ES114 (target) and MJ11 reported as the amount of target CFUs recovered after 24 hour (log CFU/mL). MJ11 genotype is shown on the x-axis: wild-type (WT), *tssF* mutant (*tssF-*), *tasL* mutant (*tasL-*), and *tasL* and *flgK* mutants simultaneously (*tasL-flgK-*). Letters indicate significantly different target recovery values between treatments (one-way analysis of variance (ANOVA) with Tukey’s multiple comparison test; *P<*0.0001). Each experiment was performed four times with three biological reps (n=12). (C) Model for how coculture of *tasL* and *flgK* mutants can complement aggregation and target killing in hydrogel.

Given that aggregation ability is directly correlated with killing in hydrogel, we predicted that the *flgK* and *tasL* coculture would also have enhanced killing ability, relative to these strains acting alone. To test this prediction, we performed coincubation assays between ES114 (target) and MJ11 (inhibitor) strains in hydrogel. The number of recovered ES114 CFUs was significantly higher with the T6SS2 structural mutant (*tssF-*), *tasL* mutant, and flagella mutant (*flgK-*) compared to the wild type (Fig 5B), which is consistent with previous observations that T6SS2, TasL, and flagella are required for killing in hydrogel. When ES114 was incubated with the *tasL* and *flgK* mutants simultaneously, however, recovered ES114 CFUs were significantly lower than with the *tasL* and *flgK* mutants alone, but significantly higher than the wild type (Fig 5B), suggesting that killing was partially restored. This observation supports the hypothesis that flagella-dependent aggregation factors and TasL may act in concert to facilitate cell-cell contact and thus killing (Fig 5C).

### Flagella promote transcriptional changes of symbiotically relevant genes

Given our previous observations that environmental viscosity primes *V. fischeri* cells for T6SS killing within host-like environments (12), we predicted that the viscosity signal may prime cells for other host-relevant functions. Therefore, we searched the transcriptomes for genes with previously characterized, host-relevant functions that showed *flgK*-dependent increased transcription in hydrogel relative to liquid media. This analysis identified genes relating to motility, chemotaxis, carbon and iron transport, and nitric oxide detoxification (Fig 6).

**Figure 6.**
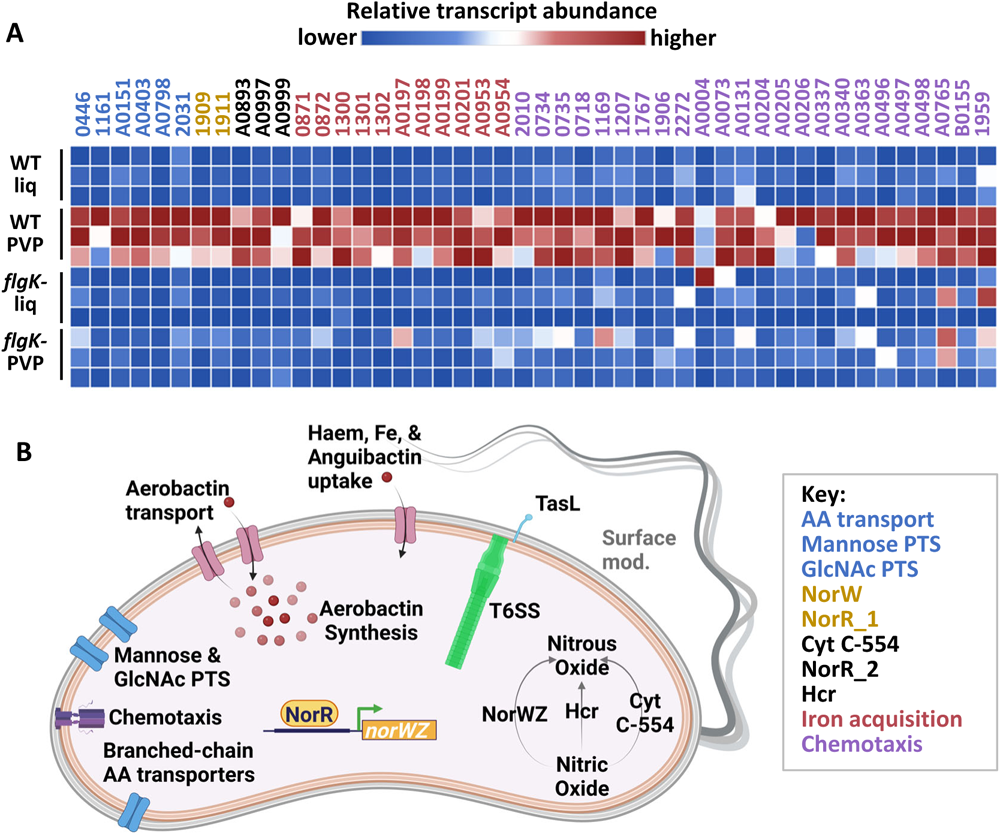
Flagella coordinates expression of host colonization factors with T6SS. Biological triplicate transcript abundances (transcripts per 1000 cells) for indicated VFMJ11 gene IDs relating to PTS and branched-chain amino acid uptake, aerobactin synthesis and transport, iron acquisition pathways, nitric oxide detoxification, and chemotaxis-related proteins. Treatments include the wildtype MJ11 grown in either low viscosity LBS medium (WT liq) or high-viscosity LBS medium (WT PVP), and the MJ11 *flgK* mutant grown in either low viscosity (*flgK-* liq) or high viscosity (*flgK-* PVP). All genes showed significantly higher expression in WT in hydrogel (>3-fold) compared to WT liquid, which was dependent on *flgK*. (B) Diagram of the contribution of flagella to coordinate expression of competition and host colonization factors in hydrogel for *V. fischeri* MJ11. T6SS and TasL expression is independent of *flgK* (Fig 4), and all other functions are upregulated in hydrogel in a *flgK*-dependent manner (A). Molecular structures are not displayed to scale. Panel A was made with Morpheus (http://software.broadinstitute.org/Morpheus) and panel B was made with Biorender.com.

Motility and chemotaxis are required for host colonization in *V. fischeri* (32–35). We observed that 18 out of 54 flagella genes had significantly higher transcript abundance in wild-type hydrogel samples relative to wild-type liquid and *flgK* mutant samples (Fig S7). These data are consistent with previously published proteomes where flagella-related proteins, such as filaments, were more abundant in cells grown in hydrogel (12), and suggest flagella may be required to enhance their own expression during habitat transition. We also observed a *flgK*-dependent increase in transcript abundance in hydrogel for chemotaxis-related genes including methyl-accepting chemotaxis proteins and corresponding regulatory proteins CheR and CheW (Fig 6A, Table S2).

Previous work also uncovered some of the nutrients that are available or limiting within the host light organ, including the presence of mannose (36), N-acetylglucosamine (37), amino acids (18), and haem, which can serve as an iron source (38). We found the transcripts for N-acetylglucosamine and mannose-specific PTS genes, as well as branched chain amino acid transporter genes were expressed up to 10-fold higher in wild-type hydrogel samples compared to other treatments (Fig 6A). Furthermore, transcript abundance for synthesis of the aerobactin siderophore and transporter gene cluster (39, 40) as well as the haem uptake genes (38) were also highly expressed in hydrogel (Fig 6A). Because cells appear to be expressing carbon and iron acquisition pathways under conditions where they are not directly experiencing iron limitation (LBS + PVP) or the presence of these specific sugars, these results suggest that higher viscosity is a signal to prepare for the nutrient conditions that are encountered in the host.

Finally, *V. fischeri* cells encounter host-derived nitric oxide (NO) and require detoxification mechanisms to promote colonization (41–45). Interestingly, we found that hydrogel stimulates expression of two activators (*norR_1* and *norR_2*) of NO detoxification pathways in other organisms: cytochrome c-554 and nitric oxide reductase (*hcp/hcr*) (46–48) (Fig 6B). However, expression of *V. fischeri* genes that are known to be induced by NO (ex. *aox*) were not elevated in our transcriptomes (45, 49) (Table S2), indicating that the cells are not experiencing NO stress but rather preparing for it, in response to transition to high viscosity. Taken together, these transcriptional effects suggest that flagella are necessary to coordinately activate expression of key competition and colonization factors in *V. fischeri*.

## Discussion

This work combines a genome-wide screen with transcriptomics to show the key role of flagella in facilitating competitive interactions in a liquid environment. Through quantitative transcriptomics, we determined that *V. fischeri* dramatically alters its cellular mRNA content by elevating global transcription during simulated habitat transition between free-living and host-associated lifestyles. Our evidence suggests flagella serve as a possible mechanosensor to detect and respond to changes in environmental viscosity during habitat transition in two important ways: 1) activating expression of surface modification genes required for aggregation necessary for T6SS-mediated killing, and ii) modulating expression of other symbiotically-relevant genes.

Direct and indirect evidence for flagella as a mechanosensor has been described for *E. coli*, *Proteus*, *Pseudomonas*, *Vibrio cholerae* and *Vibrio parahaemolyticus*, (50–52). For these model organisms, the flagella sense the physical environment to initiate highly regulated, sequential processes that direct coordinated cellular behaviors such as biofilm and swarming motility. In the case of *V. parahaemolyticus*, the closest relative of *V. fischeri* in this group, mechanical inhibition of the flagella rotation due to high viscosities, surfaces, or functional disruption (via phenamil or mutation), can trigger swarmer cell differentiation (53, 54), elicit upregulation of colonization and virulence factors (55), as well as increased expression and secretion of the T6SS protein Hcp1 (56). Similarly, our data support a model whereby *V. fischeri* flagella are required to elicit a transcriptional response to host-like viscosity to coordinate multiple behaviors that are required for successful competitive colonization of a host. Future work may identify the mechanism(s) by which *V. fischeri* flagella detect changes in environmental viscosity, as well as additional sensors that activate the T6SS weapon.

Our findings also provide a possible explanation for why T6SS2 in *V. fischeri* requires the Sigma 54-dependent transcriptional regulator VasH to fully activate T6SS2. Previous work showed the T6SS2-encoding *V. fischeri* strain FQ-A001 requires *rpoN*, which encodes the Sigma-54 cofactor, for both T6SS2 function and motility (57). Our data suggest that T6SS2 function and motility may be coordinately regulated through the requirement of a common sigma factor to facilitate cell surface modifications that promote aggregation, which is required for T6SS-mediated killing in liquid environments. Similar mechanisms of coordinate regulation have been observed in other taxa where sigma factors play important roles in coregulating virulence factors such as T6SS and flagellar motility (58–61).

In conclusion, this work identifies novel factors that promote contact-dependent competition in a viscous liquid and reveals how the physical environment influences expression of symbiotically-relevant genes. Based on these and previous findings we propose an updated model for the role of T6SS in symbiosis establishment between *V. fischeri* and its host (Fig 7). Juvenile squid hatch with an aposymbiotic light organ that is colonized by free-living *V. fischeri* from the water column. Aposymbiotic squid produce highly-viscous extracellular mucus that coats the light organ and promotes aggregation of *V. fischeri* on the light organ surface. Cells then migrate into the light organ through pores that lead to six physically separated crypt spaces containing a highly-viscous lumen. Our findings suggest that the *V. fischeri* flagella are required for transcriptional and behavioral responses to changes in ambient viscosity associated with symbiosis establishment. This transition to a highly-viscous host environment coordinately activates expression of functions to promote the survival and growth of *V. fischeri* cells, as well as compete with other genotypes that attempt to colonize the same crypt. In the event that a crypt is colonized by incompatible *V. fischeri* strains, surface-modification factors and TasL allows T6SS2+ strains to make contact with and kill competitor cells.

**Figure 7.**
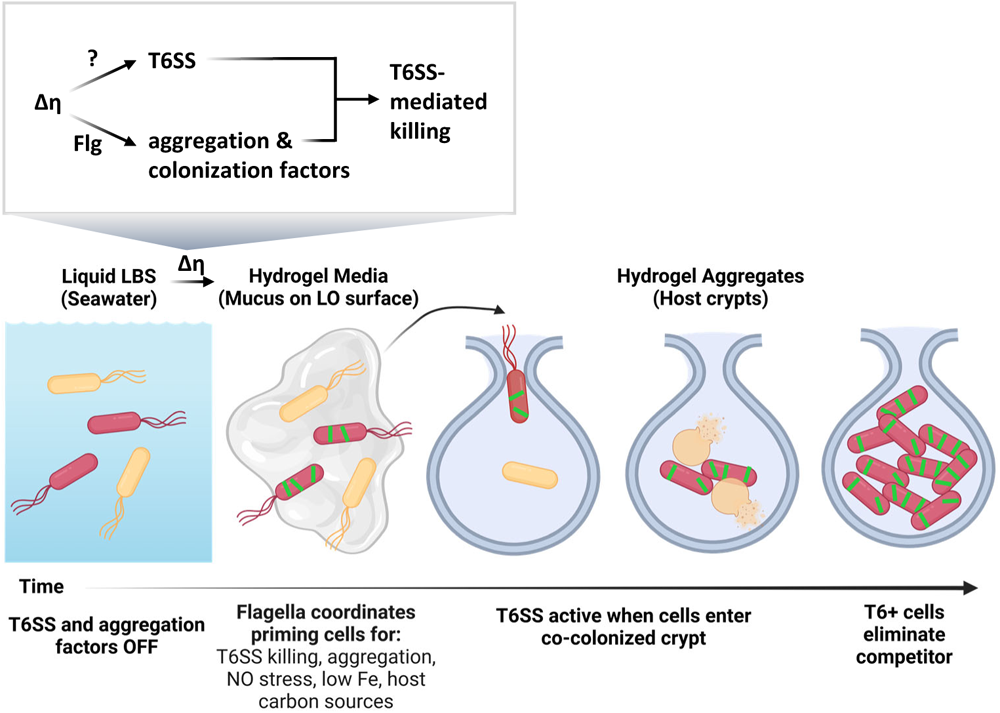
Model for how flagella primes cells for T6SS-mediated competition within the host. Bacterial cell colors indicate incompatible *V. fischeri* strains without (yellow) or with (red) a T6SS2 gene cluster. From left to right: free-living cells begin in a lower-viscosity liquid environment before associating with viscous, host-derived mucus on the surface of the light organ. Flagella sense the change in environmental viscosity (Δƞ) to activate pathways associated with aggregation, iron and carbon acquisition, and nitric oxide detoxification. The genetic determinants of T6SS activation during habitat transition remains unknown and is independent of flagella. Cells then migrate into viscous lumen of the crypt space where T6+ cells kill competitors and proliferate to dominate the crypt space. Figure made with Biorender.com.

## Methods

See text S1 for additional methods and experimental details, specifically media and growth conditions, strains and plasmids, details about the screening transposon mutants, coincubation assays, motility assays, and fluorescence microscopy.

### Transposon mutagenesis and screen for conditional mutants

Conjugations were set up between *V. fischeri* strain MJ11 (23) and the mini-Tn5-ermR transposon vector pEVS170, which was maintained in *E. coli* strain RHO3, a DAP auxotroph (62). After a 24-h incubation period on Luria Bertani with added Salt (LBS) agar supplemented with DAP at 24°C, mating spots were resuspended in 100 μl LBS medium and spread plated onto LBS agar medium containing erythromycin (erm) (see supplemental text for more details). A total of 7,300 colonies were picked and screened for the ability to kill *V. fischeri* strain ES114 in hydrogel (LBS broth with 5% w/vol PVP) yet not on LBS agar plates.

High throughput coincubation assays in hydrogel were performed for each mutant and a GFP-tagged target strain as described previously (11). Strains were mixed in a 9:1 (MJ11 tn5 mutant:ES114 pVSV102) ratio and incubated standing in LBS hydrogel for 24 hours in a 96-well plate at 24°C. Luminescence, fluorescence, and absorbance values collected by a Tecan Plate reader were used to assess the killing ability of each mutant. Only mutants from wells with bright luminescence (indicating MJ11 grew) and GFP fluorescence (indicating the target grew) values comparable to those from the control containing ES114 pVSV102 and MJ11 *tssF_2* mutant strain LAS04 were selected for further study. Of the 7,300 mutants screened 222 did not kill ES114 in hydrogel. We validated this phenotype by performing coincubation assays in hydrogel and quantifying CFUs of target strain (12), and moved forward with 45 mutants. We then performed agar plate coincubation assays, as described in (5, 63) between each mutant and GFP-tagged ES114 to determine whether each mutant prevented the growth of ES114 when contact was forced. 20 of these mutants prevented the growth of ES114 on agar plates.

### Mapping transposon insertion sites

To identify where the transposon inserted in each mutant, we performed cloning, as described in (64), and inverse PCR (iPCR). For all transposon mutants, chromosomal DNA was digested with the HhaI restriction enzyme, fragments were self-ligated with T4 DNA ligase. For cloning, the resulting circularized DNA was transformed into *E. coli* DH5⍺λpir and plated onto BHI erm to select for transformants containing the Erm resistance gene within the transposon. Plasmid minipreps were performed to purify the circularized transposon and flanking DNA and Sanger sequencing was used to sequence across the transposon-chromosome junction using primer M13F. For iPCR, the circularized gDNA from T4 DNA ligase reactions was used as the template for PCR amplification with primers specific for the transposon: AS1100 (TACATTGAGCAACTGACTGAAATGCC) and EXT170 (GCACTGAGAAGCCCTTAGAGCC). These primers sequence outward from the transposon, around the piece of circularized DNA, such that the resulting PCR product is a combination of the transposon and the disrupted gene. iPCR products were cleaned with a Zymo Clean/Concentrate kit followed by Sanger sequencing.

### Genome resequencing

LAS22A10 gDNA was extracted using a Zymogen Bacterial and Fungi DNA extraction kit, quantified with a QuantIT kit, and sent for Illumina sequencing to SeqCenter LLC (Pittsburgh, PA). Using Geneious software, after the reads were trimmed, error corrected, duplicate reads removed, they were mapped against the MJ11 reference genome, and mutations in the LAS22A10 reads were identified using the Single Nucleotide Polymorphism (SNPs) feature in Geneious, with default settings and 100% variant frequency. Mutation locations were confirmed using NCBI Blast.

### Natural Transformation

To construct strain MP101, MJ11 pLostfox-Kan was prepared as previously described ((26), supplemental text S1) and transformed with gDNA extracted from LAS22A10 (*flgK*:tn5). Transformants were screened for erm resistance and kan sensitivity to ensure transfer of the transposon insertion and loss of the pLostfox-Kan plasmid.

### Transcriptome collection and data analysis

Overnight cultures of MJ11 were subcultured into fresh LBS liquid or hydrogel and grown for 12 hours at 24°C. To dilute the PVP and stop transcription, 500 μl of each sample was added to a pre-chilled tube containing 3 mL LBS medium and 1 mL stop solution (5% v/v phenol pH 4.3 and 95% ethanol v/v). Cells were pelleted through centrifugation (4k rpm for 15 minutes at 4°C), washed with 500 µl LBS, and pellets were frozen overnight at −80°C. CFUs were quantified for each sample (three technical replicates per sample) prior to pelleting for downstream calculations. RNA was then extracted using the MirVana kit following the manufactorer’s protocol (Thermo Fischer Scientific, Waltham, MA). *Saccharolobus solfataricus* P2 RNAs were used as internal RNA standards (IS), which were added after adding lysis buffer and prior to adding the miRNA homogenate additive; see Text S1 for IS details. Residual DNA was removed via the Turbo DNA-free kit (Invitrogen, Carlsbad, CA). cDNA library prep and sequencing were performed at the University of North Carolina (UNC) High-Throughput Sequencing Facility (HTSF) with the HiSeq 4000 platform (single-end 50-bp reads). Three biological replicates for each strain genotype and treatment were sequenced (n=12). Quality scores were calculated using FastQC; low-quality sequences (average quality score <20 across 5 bp) were removed using Trimmomatic (65). rRNA and IS RNA was counted and removed from the dataset with BLAST. Reads were mapped to the MJ11 genome using BowTie2 (66) and counted using HTSeq (67). Methods described in (27, 28) were used to quantify MJ11 mRNA within each sample. A recovery factor for each IS within a sample was calculated by taking the amount of individual ISadded and dividing by the number of IS transcripts recovered. Recovery factors for the X IS were averaged for each sample. Read counts for each gene were then multiplied by the average recovery factor for each sample and then divided by the average number of CFUs (cells) for each sample to calculate the total number of transcripts per cell for each sample (See Table S2 for calculations).

Differential expression analysis was performed as described previously (12, 68). This involved a contingency table was generated by comparing average transcript per CFU abundance between treatments using a Student’s t-test corrected for multiple comparisons using the false discovery rate (FDR). Volcano plots were generated by graphing the negative log_10_ q-value and log_2_ fold change between treatments. Because zero values cannot be log-transformed for differential expression analysis, zero values were set to the limit of detection for a given gene, which was one read per gene. Only transcripts with a log_10_q-value ≤ 0.05 and a log_2_ fold change ≥ |1| are considered statistically significantly differentially expressed (DE). Hierarchical clustering analysis was performed using the heatmap function in R; transcript per 1000 cell values were not scaled. The vegan package in R was used to perform principal coordinates analysis (PCoA) using the Bray-Curtis dissimilarity index, PerMANOVA using the Adonis function. (69). Dispersion was determined using the ‘permutest.betadisper’ function to ensure there were no significant differences in dispersion between the groups being compared. Transcriptome sequences were submitted under Bioproject number PRJNA1013100.

## Acknowledgements and Funding Sources

We thank Peggy Cotter, Andreas Teske, Barbara MacGregor, Manuel Kleiner, Anne Dunn, and Andrea Suria for helpful discussions, and the UNC Microscopy Services Laboratory for SEM sample preparation and imaging assistance. Work in the lab of Alecia Septer was supported by NIGMS grant R35 GM137886. Lauren Speare was supported by a UNC Dissertation Completion Fellowship and as a Simons Foundation Awardee of the Life Sciences Research Foundation. We declare no conflict of interest.

---

**Table S1.**
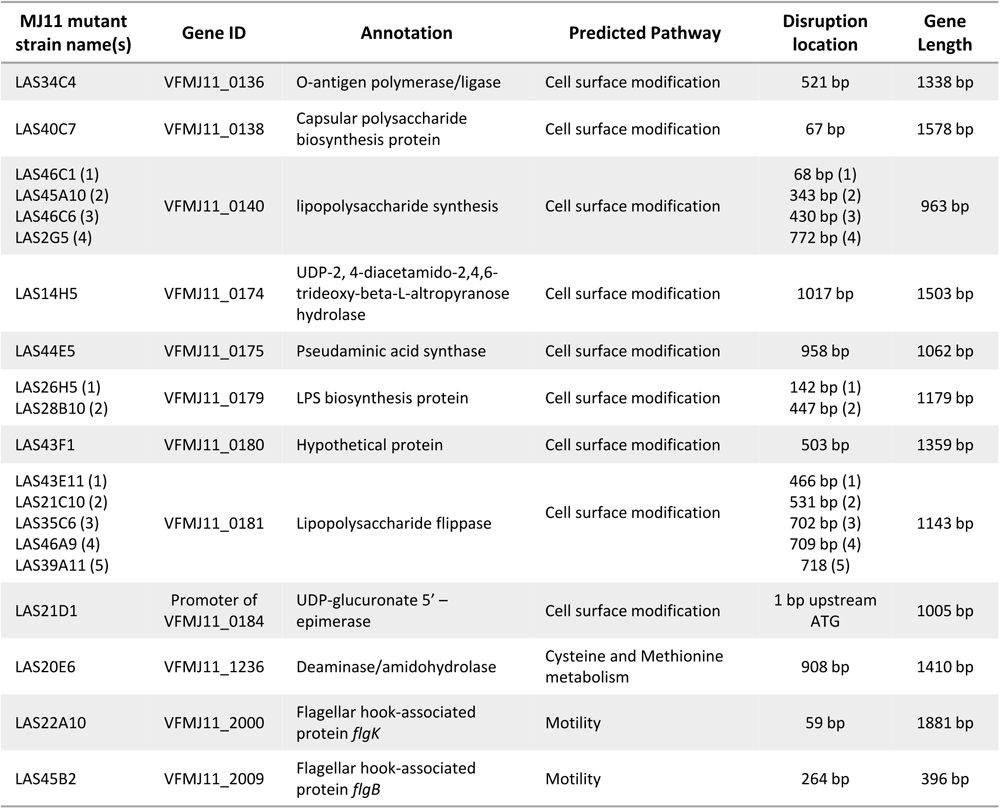
Location of each transposon insertion in the MJ11 genome. Annotation and predicted pathways provided by KEGG.

**Figure S1.**
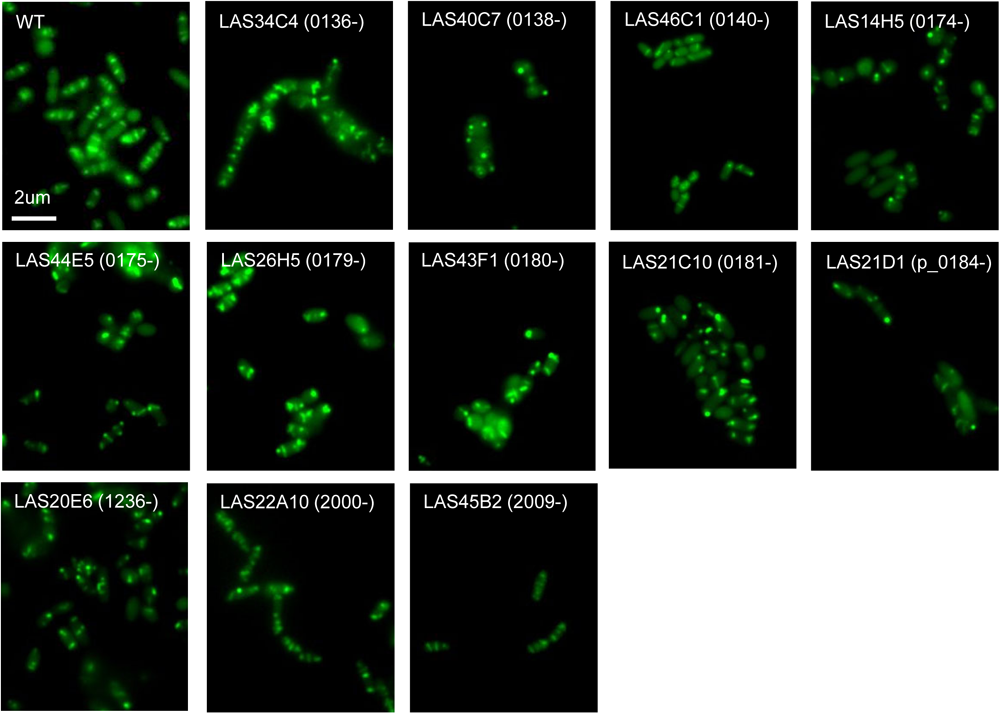
Conditional mutants can assemble T6SS2 sheaths. Representative fluorescence microscopy images of MJ11 strains harboring an IPTG-inducible VipA_2-GFP expression vector. Cultures were incubated in hydrogel supplemented with 1.0 mM IPTG; scale bar = 2 um. Strain genotype is indicated as follows: transposon inserted into VFMJ11_0136 (0136-). WT indicates wild-type. Each experiment was performed at least twice with two biological replicates and five fields of view per replicate (n=20).

**Figure S2.**
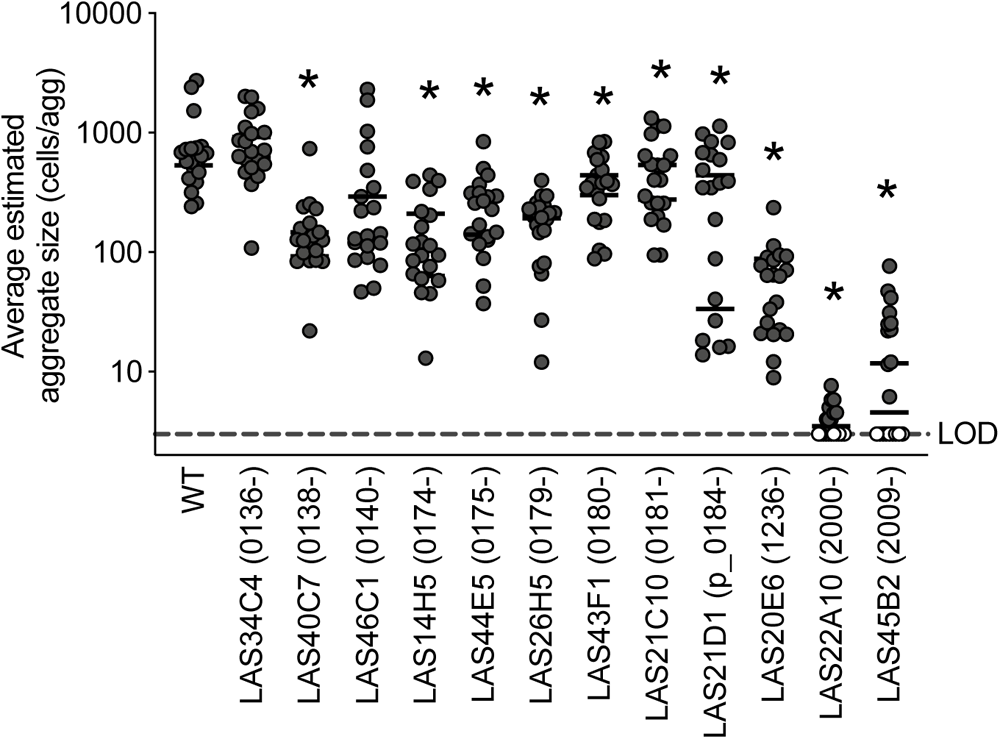
Conditional mutants show reduced aggregation in hydrogel. Average estimated aggregate size (shown as cells per aggregate) for MJ11 strains. Strain names and genotypes are shown on the x-axis. LOD indicates the limit of detection (3 cells touching one another) and is shown by white symbols. Asterisks indicate significantly lower average estimated aggregate size than the wild type (WT) (two-way analysis of variance [ANOVA], *p<0.003*). Each experiment was performed twice with two biological replicates and five fields of view (n=20).

**Figure S3.**
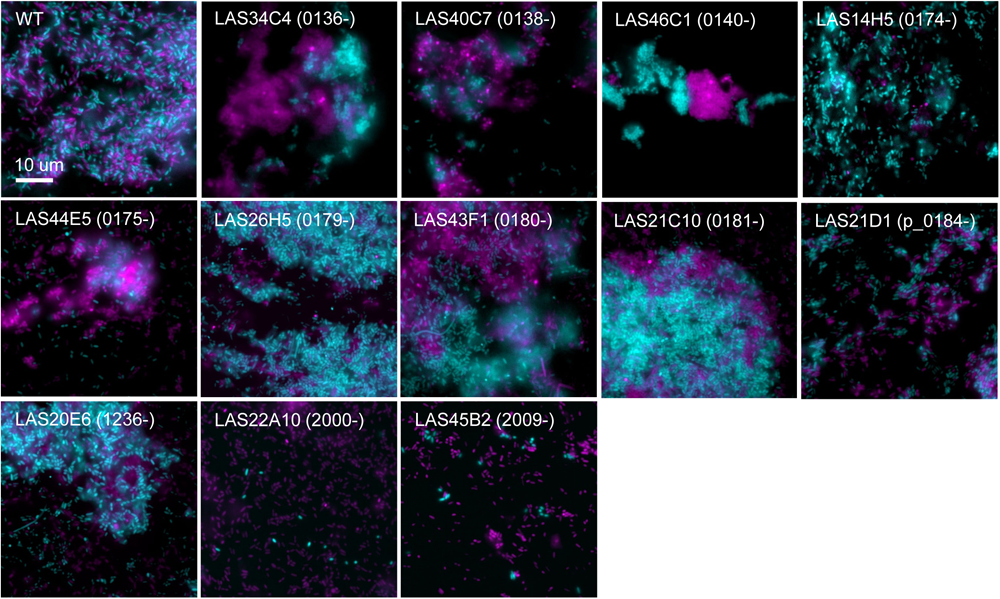
The majority of conditional mutants form mixed aggregates. Representative single-cell fluorescent microscopy images of differentially-tagged but otherwise isogenic MJ11 co-cultures incubated in hydrogel for 12 hours; scale bar = 10 um. Each experiment was repeated twice with two biological replicates and 3 fields of view (n=12).

**Figure S4.**
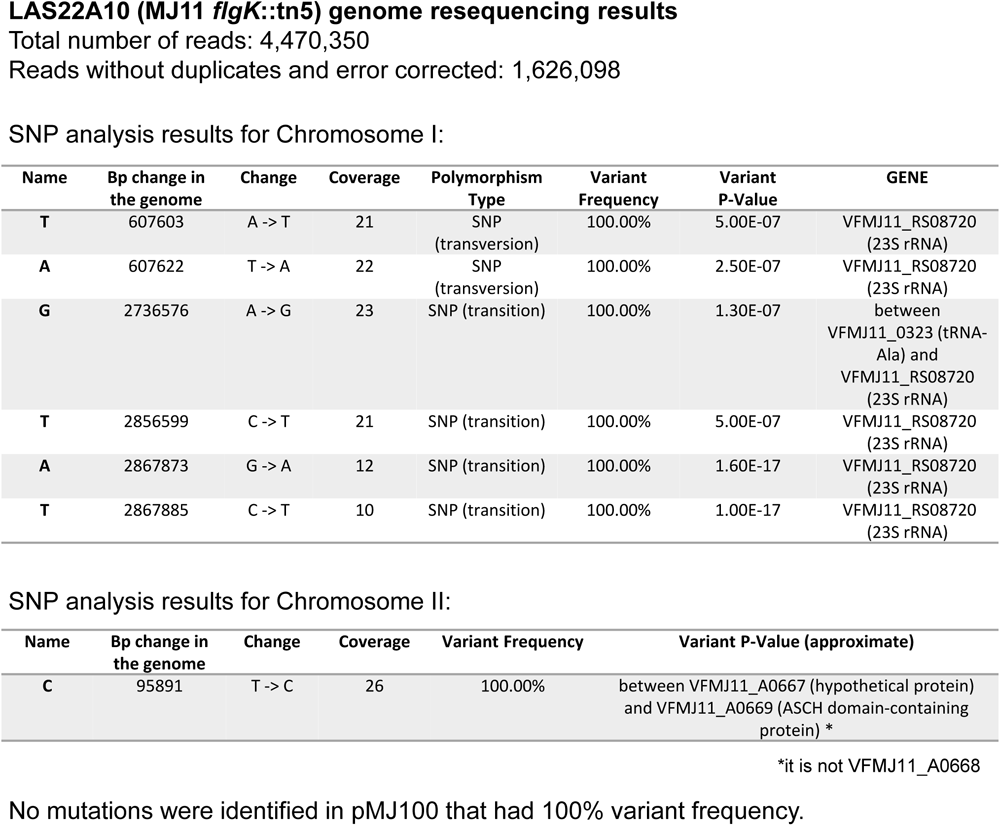
LAS22A10 resequencing results. Trimmed, error corrected, and duplicate removed reads for LAS22A10 (flgK::tn5) were mapped to the MJ11 reference genome using Geneious software. A SNP analysis was performed using Geneious with default settings. Variants with 100% frequency were identified and confirm using NCBI Blast.

**Figure S5.**
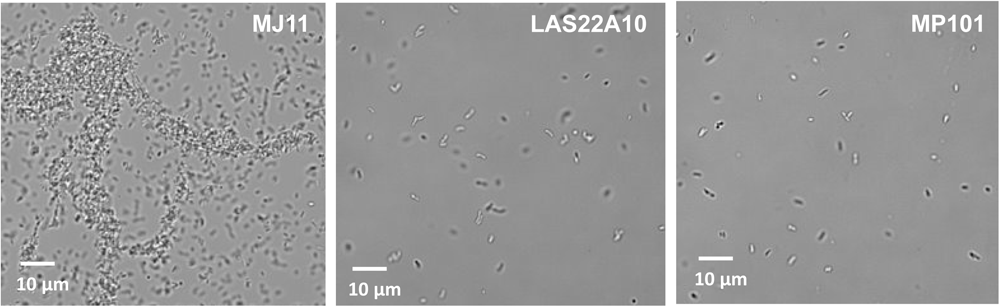
Aggregation phenotypes for original and remade *flgK*::tn5 mutants. *V. fischeri* strains were grown on LBS agar with the appropriate antibiotic. Three milliliters of liquid LBS with PVP was inoculated and grown at room temperature without shaking. When the cultures reached an OD_600_ of 0.5 to 1.0, two microliters of the culture were placed on a glass slide with coverslip and visualized for using a Nikon Ti2 microscope with NIS-Elements software. Each experiment was performed three times with 2 biological replicates and 5 fields of view.

**Figure S6.**
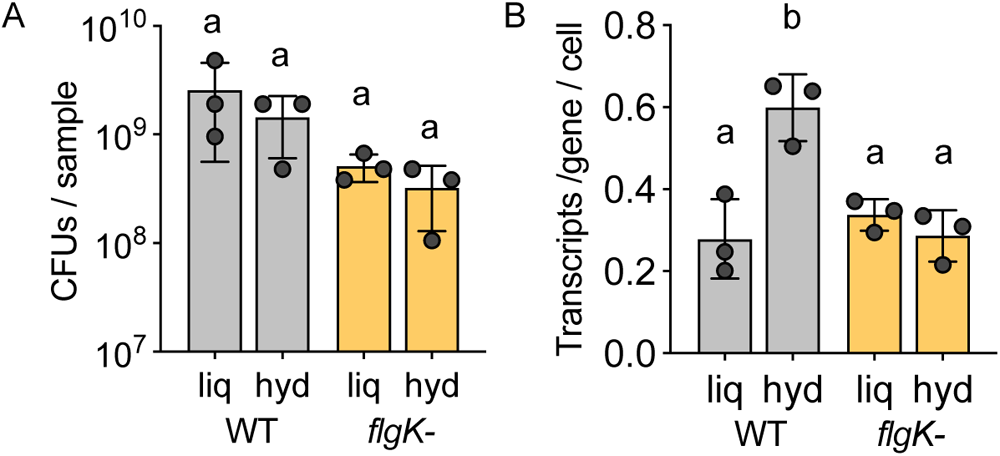
Statistics for transcriptomes. (A) Number of colony forming units (CFUs) per cell and (B) number of transcripts per gene per cell for MJ11 transcriptomes for the wild-type (WT) or *flgK* mutant (*flgK-*) incubated in liquid LBS (liq) or hydrogel (hyd). Each circle represents an individual sample (transcriptome). Letters indicate significantly different values between sampling groups (two-way analysis of variance [ANOVA] with Šídák’s multiple comparison post-test, *p<0.02*).

**Figure S7.**
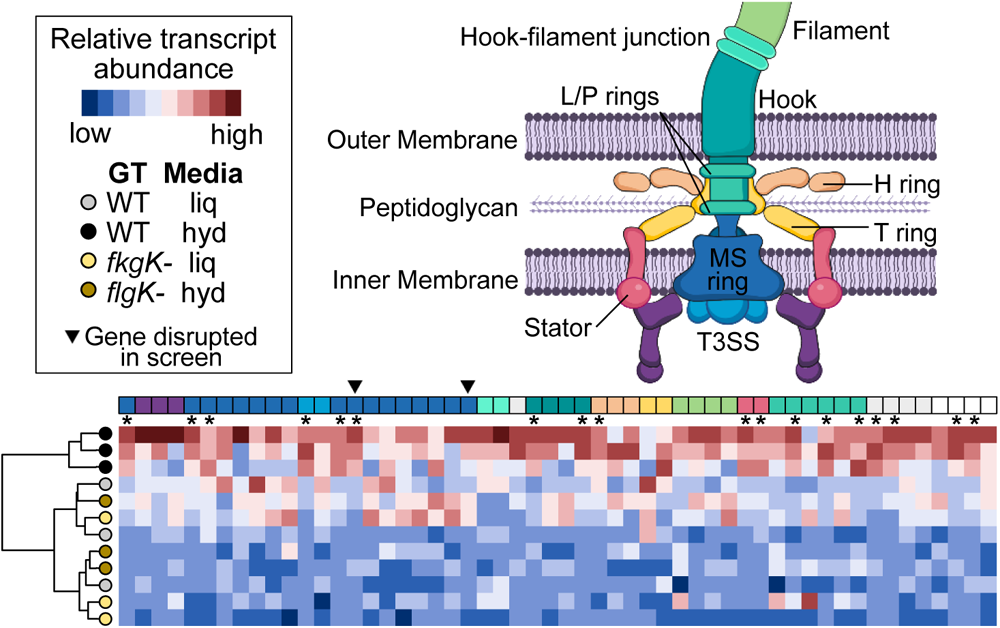
Flagella promotes elevated expression of flagella-related genes in hydrogel. Hierarchical clustering analysis and heatmap displaying relative transcript abundance for flagellar-associated genes. Circle color indicates the strain genotype and media condition: light gray (wild-type [WT] in liquid [liq]), dark gray (wild-type in hydrogel [hyd]), light yellow (*flgK* mutant [*flgK-*] in liquid), and dark yellow (*flgK* mutant in hydrogel). Transcript abundance is shown as relative transcript abundance were values are scaled across each gene (column); blue indicates relatively low transcript abundance while red indicates relatively high abundance. Squares above each column (gene) indicate its known or predicted function based on schematic shown at the top of the figure. Asterisks indicate expression of a gene was significantly higher for wild-type hydrogel samples compared to all other samples (one-way ANOVA, *p<0.03*).

