## Supplementary material for "Flagella are required to coordinately activate competition and host colonization factors in response to a mechanical signal": Text S1

**Supplemental methods**

**Media and growth conditions.** *V. fischeri* strains were grown in LBS media (1) that was not supplemented with polyvinylpyrrolidone (PVP) or was supplemented with 5% w/v PVP (hydrogel) (2) at 24°C and *E. coli* strains were grown in either LB medium or Brain Heart Infusion (3) at 37°C that was supplemented with 400 μg/ml diaminopimelic acid (DAP) for *E. coli* Rho3. Antibiotic selection for *V. fischeri* strains are as follows: chloramphenicol (2 μg/ml), erythromycin (5 μg/ml), and kanamycin (100 μg/ml) (4). Antibiotic selection for *E. coli* strains are as follows: chloramphenicol (20 μg/ml) and kanamycin (40 μg/ml) in LB, and erythromycin (150 μg/ml) in BHI. Plasmids pEVS102 and pEVS208 were maintained in *E. coli* DH5⍺λpir (4); plasmid pEVS104 (5) was maintained in *E. coli CC118λpir* (6) and plasmid pEVS170 was maintained in *E. coli* Rh03 (7).

**Additional strains and plasmids used in this study.**

| **Strain** | **Relevant characteristics** | **Reference** |
| --- | --- | --- |
| ES114 | *V. fischeri;* isolated from *Euprymna scolopes* light organ | Boettcher and Ruby 1994 |
| MJ11 | *V. fischeri;* isolated from *Moncentris japonica* light organ | Ruby and Nealson 1976 |
| LAS04 | MJ11 with a disruption in *tssF_2* (Erm^R^) | Speare *et al.,* 2021 |
| LAS015 | MJ11 with a disruption in *tasL* (Erm^R^) | Speare *et al.,* 2022 |
| MP101 | MJ11 *flgK*::tn5 remade from LAS22A10 gDNA | This study |
| *E. coli* CC118λpir | *E. coli; Δ(ara-leu) araD Δlac74 galE galK phoA20 thi-1 rpsE rpsB argE*(Am) *recA λpir* | Herrero *et al.,* 1990 |
| *E. coli* DH5⍺λpir | *λpir derivative of E. coli; F’/endA1 hsdR17 glnV44 thi-1 recA1 gyrA relA1* Δ*(lacIZYAargF)*  *U169deoR(f80dlacI* Δ(*lacZ)M15)* | Dunn *et al.,* 2005 |
| *E. coli* Rho3 | *E. coli;* SM10(λ*pir*) Δ*asd*::*FRT* Δ*aphA*::*FRT; Erm^R^* | López *et al.,* 2009 |
| **Plasmids** | **Relevant characteristics** | **Reference** |
| pEVS104 | conjugative helper, *oriV_R6k_*_γ_, *oriT, Kn^R^* | Stabb and Ruby 2002 |
| pVSV102 | *gfp+, oriV_R6k_*_γ_, *oriV_pES213_, oriT, Kn^R^* | Dunn *et al.,* 2006 |
| pVSV208 | *dsRed+,* *oriV_R6k_*_γ_, *oriV_pES213_, oriT, Cm^R^* | “” |
| pEVS170 | mini-Tn5 delivery vector maintained in Rho3, *oriV_R6k_*_γ_, oriT, Erm^R^, | Lyell *et al.,* 2008 |
| pLostfox-Kan | *tfoX* expression vector, *oriT*, *f1 ori*, Kan^R^ | Brooks *et al.,* 2014 |
| pSNS119 | *vipA_2-gfp* fusion vector*; oriVR6Kγ, oriVpES213, oriT, Kn^R^* | Speare *et al.,* 2018 |

**Transposon screen for conditional mutants**.

To determine whether transposons disrupted genes that are conditionally required for T6SS2 killing, we performed high throughput coincubation assays in LBS hydrogel between each mutant and a target strain (ES114) derived from methods described previously (8). Transposon mutant colonies were grown overnight in 96-well plates containing 200 μl LBS Erm liquid medium and overnight cultures of GFP-tagged ES114 were grown in LBS kanamycin liquid media. Each transposon mutant was mixed in a 9:1 ratio (by volume) MJ11:ES114 and each mixture was spotted into a new 96-well plate containing 200 μl hydrogel and incubated at 24°C without shaking. At 24 h the luminescence, fluorescence, and absorbance of each well was measured to determine whether each strain grew during the experiment. Four controls were included in each 96-well plate and contained: only media to account for background luminescence and fluorescence values from the hydrogel, the target strain alone (representing expected luminescence if the mutant was unable to grow), the target with MJ11 wild-type (a killing control), and the target and a MJ11 *tssF* disruption mutant (a no-killing control). Luminescence and fluorescence values from these control wells were used to determine the outcome of coincubations with each transposon mutant. Only mutants from wells with bright luminescence and fluorescence values comparable to those from the control containing ES114 and MJ11 *tssF-* were selected for further study. Using this scaled-up version of our hydrogel coincubation assay, we screened 7,300 mutants and identified 222 strains with mutations that no longer inhibit the growth of ES114. We chose to move forward with 45 mutants. We validated the results of our transposon screen by performing coincubation assays between each mutant and ES114 where we quantified each strain at the beginning and end of our experiment by plating for CFUs. Coincubation assays were performed as described previously (2).

We then performed agar plate coincubation assays, as described in (9, 10) between each mutant and GFP-tagged ES114 to determine whether each mutant prevented the growth of ES114 when contact was forced. Of the 45 strains screened, 21 prevented the growth of ES114 on agar plates. One of these mutants was a false positive; the effect of the mutation inhibited killing in hydrogel, however the mutant was able to kill in hydrogel during longer coincubation assays and was therefore not included in downstream experiments. We chose to move forward with the remaining 20 mutants which were unable to kill ES114 in hydrogel but still prevented the growth of ES114 on agar plates.

**Natural Transformation (for MP101 construction).** An overnight culture of the *Vibrio* *fischeri* strain MJ11 containing the plostfox-Kan plasmid was incubated in LBS with Kanamycin at 24°C with shaking. The cells were then centrifuged at 15,000 rpm for 1 minute and resuspended in 1 mL Tris-GlcNAc-kan. Thirty microliters of the cell suspension was added to 3 mL Tris-GlcNAc-Kan and incubated at 24°C to an OD_600_ of 0.2. When cells reached an OD_600_ of 0.2, 500 μL of culture was transferred to a microcentrifuge tube with 2.4 µg of LAS22A10 gDNA, which was extracted using a Zymo Bacterial and Fungal DNA kit. The cells were vortexed briefly and left at room temperature for 30 minutes. Cells were then transferred into a glass tube with 1 mL LBS liquid and incubated at 24°C overnight with shaking. One hundred microliters of the overnight culture were plated onto LBS containing Erythromycin to select for the flgK::tn5 transposon insertion and incubated for 18 hours at 24°C. Four colonies picked and restreaked on LBS Erythromycin plates. The next day the cultures were verified by growing on selective LBS Kanamycin and LBS Erythromycin plates at 24°C overnight, to ensure the plostfox-Kan plasmid had been lost and the erm-resistant transposon insertion retained.

**Coincubation assays with the *tasL* mutant.** ES114 pVSV102, MJ11 wildtype, MJ11 *vasA*_2 (LAS04), MJ11 *tasL* mutant (LAS015), and MJ11 *flgK*::tn5 (LAS22A10) were streaked from glycerol stocks onto LBS medium (supplemented with appropriate antibiotic) and incubated at 24°C overnight. Individual colonies were restreaked onto LBS plates and incubated overnight at 24°C. Cells were resuspended in LBS liquid medium and normalized to an OD of 1.0. A 12-well plate was filled with LBS with 5% PVP and 2 ul of ES114 pVSV102 was added to each well, along with 20 ul of the MJ11-derived strain. Plates were incubated at room temperature for ~24 hours. Serial dilutions of each coincubation were plated onto LBS Kan plates to select for and quantify the remaining ES114 pVSV102 target strain.

**VipA_2 sheath visualization and quantification.** Visualization of VipA_2-GFP sheaths was performed as described previously (2, 9). Single-cell fluorescence microscopy was performed by visualizing cultures of MJ11 harboring the IPTG-inducible VipA_2-GFP (TssC) expression vector pSNS119 (9). Cultures were grown overnight in liquid LBS supplemented kanamycin and 0.5 mM isopropyl-β-D-1-thiogalactopyranoside (IPTG), then diluted 1:100 into fresh hydrogel media with the same supplements and incubated for 2.5 hours. 3 μl of each culture was spotted onto a glass slide, covered with a cover slip, and imaged using an Olympus BX51 microscope outfitted with a Hammatsu C8484-03G01 camera and a 100X/1.30 Oil Ph3 objective lens. Images were analyzed using Fiji.

**Motility Assays.** Overnight cultures of each strain were diluted into fresh LBS media supplemented with Erm and grown to an OD_600_ of 0.4 and 10 μl of each culture was spotted onto LBS plates with 0.3% agar. Spot diameter was measured every six hours for 18 hours and results are reported as percent of wild-type. We chose to use LBS rather than SWT media, which is most commonly used to assess *V. fischeri* motility (11), to understand how each mutation affected swimming in LBS-derived medias, in which all other experiments were performed, and avoid any confounding results which could be linked to the salts in instant ocean.

**Aggregation visualization and quantification.** Aggregates were visualized and quantified as described previously (2, 12). Briefly, overnight cultures of *V. fischeri* containing either pVSV102 (GFP) or pVSV208 (DsRed) grown in LBS supplemented with either kanamycin or chloramphenicol, respectively, were normalized to an OD_600_ of 1.0 and 10 μl spotted into 1 mL hydrogel. In the case of differentially tagged yet otherwise isogenic cultures, strains were mixed in a 1:1 ratio after normalization and prior to being inoculated into hydrogel. Cultures then incubated in hydrogel for 12 hours, not shaking. 3 μl of each culture was spotted onto a glass slide and imaged with a 60 /1.3 numerical aperture oil Ph3 lens objective. Images were captured with an Olympus BX51 microscope outfitted with a Hamamatsu C8484-03G01 camera using MetaMorph software. Estimated average aggregate size was calculated by using the ”image/adjust/threshold” and “analyze particles” commands as described previously (8, 12). For coaggregation images using *flgK* and *tasL* mutants (Fig 5A), a single colony of each differentially tagged strain was used to inoculate a LBS PVP culture and grown to OD between 0.5 and 1.0. 2 µl of the coculture were then added to a glass slide and imaged using a Ti2 Nikon inverted fluorescence microscope and NIS-Elements acquisition software and Orca Fusion C1440 camera. Red and green fluorescence images were overlaid in NIS-Elements and assigned color using FIJI. Each experiment was performed three times with 2 biological replicates and 5 fields of view.

**SEM.** Overnight cultures of each strain were diluted into fresh liquid LBS media and grown for 12 h. Following the 12 h incubation period, cells were fixed for microscopy by centrifuging the culture for 1 minute at 3,000 rpm and resuspending the pellet in 4% PFA in marine PBS (standard PBS solution with NaCl added to a salinity of 20 psu to prevent lysing of marine cells). Fixed samples were prepared for SEM by the UNC Microscopy Services Laboratory and imaged on a Zeiss Supra 25 FESEM.

**Internal Standard Details**. *Saccharolobus solfataricus* P2 transcripts were used as internal RNA standards (IS). ISs were grouped into three groups and added to each sample at the following number of copies: Group 1 - locus tags SSO1129 (8.69 x 10^8^copies), SSO1378 (1.04 x 10^8^ copies), and SSO1123 (8.38 x 10^6^ copies); Group 2 – locus tags SSO1692 (8.42 x 10^8^ copies), SSO1328 (1.77 x 10^8^ copies), and SSO1806 (1.01 x 10^7^ copies); and Group 3 – locus tags SSO1273 (6.85 x 10^8^ copies), SSO1886 (1.22 x 10^8^ copies), and SSO3006 (3.54 x 10^6^ copies). ISs were added after adding lysis buffer and prior to adding the miRNA homogenate additive. DNA was removed via the Turbo DNA-free kit (Invitrogen, Carlsbad, CA). cDNA sample libraries were prepared using Tecan Genomics (NuGen) universal total RNA seq kit and sequenced at the University of North Carolina (UNC) High-Throughput Sequencing Facility (HTSF) with the HiSeq 4000 platform (single-end 50-bp reads). Three biological replicates for each strain genotype and treatment were sequenced (n=12). Quality scores were calculated for each sequence using FastQC; low-quality sequences (average quality score <20 across 5 bp) were removed using Trimmomatic (13). rRNA and IS RNA was counted and removed from the dataset with BLAST. Reads were mapped to the MJ11 genome using BowTie2 (14) and counted using HTSeq (15). To allow for comparative analyses, all genes with zero reads per gene were set to the limit of detection, which was one reads per gene. Any genes with no reads for all replicates across all treatments were not corrected for the limit of detection, and were excluded from further analysis.
